# AI-Assisted Discovery and Optimization of Small Molecule TREM2 Agonists with Functional Microglial Activity

**DOI:** 10.1101/2025.09.11.675743

**Authors:** Sungwoo Cho, Tibor Viktor Szalai, Farida El Gaamouch, Dávid Bajusz, György M Keserű, Moustafa Gabr

## Abstract

Triggering receptor expressed on myeloid cells 2 (TREM2) is a microglia-specific receptor whose loss-of-function variants increase Alzheimer’s disease (AD) risk by impairing plaque compaction, survival, and protective microglial programming. While antibody-based agonists have shown promise, their translation is hindered by poor brain penetration and high cost. Here, we report the discovery and optimization of small molecule TREM2 agonists through an AI-assisted virtual screening strategy. Deep Docking of over five million purchasable compounds identified a structurally novel hit, **T2K-014**, which engaged TREM2 with modest affinity. A SAR-by-catalog campaign led to the identification of **T2M-010** as a potent binder. **T2M-010** demonstrated favorable in vitro PK properties, including high solubility, passive BBB permeability, moderate metabolic stability, and minimal safety liabilities. Functionally, **T2M-010** activated receptor-proximal signaling, inducing SYK phosphorylation in TREM2-expressing cells, and promoted microglial phagocytosis. Together, these findings establish **T2M-010** as the most potent small molecule TREM2 binder reported to date capable of driving protective microglial responses relevant to AD.

---

Alzheimer’s disease (AD) continues to represent an urgent clinical problem, as currently approved drugs offer modest symptomatic improvement but fail to halt or significantly slow disease progression.^1^ The disorder is characterized by several pathological hallmarks: extracellular aggregates of amyloid-β (Aβ), intracellular deposits of hyperphosphorylated tau in the form of neurofibrillary tangles, and chronic activation of neuroinflammatory pathways.^2^ Within the central nervous system (CNS), inflammation is largely driven by microglia, the resident immune cells of the brain.^3^ When persistently activated, microglia release cytokines and other pro-inflammatory mediators that create a toxic microenvironment, promoting neuronal injury and contributing to the progression of AD pathology.^4^ Although many therapeutic programs have targeted neurotransmitter modulation, Aβ production and aggregation, or tau pathology, increasing attention has turned toward strategies that regulate microglial activity and inflammatory signaling as a novel avenue for intervention.^5,6^

Triggering receptor expressed on myeloid cells 2 (TREM2) is a microglia-specific receptor enriched in the CNS that signals through the adaptor DAP12/TYROBP, whose ITAM domain regulates phagocytosis, motility, proliferation, survival, and lysosomal activity.^7-10^ Rare TREM2 variants markedly increase AD risk, highlighting its central role in pathogenesis.^11,12^ In amyloid models and in R47H carriers, loss of TREM2 function impairs microglial recruitment and plaque compaction, weakens survival, and increases neuronal susceptibility to Aβ toxicity.^11,12^ Transcriptomic studies show TREM2 deficiency blocks adoption of a protective disease-associated state, with neuritic degeneration consistently exacerbated despite variable effects on total amyloid load.^11^ Conversely, TREM2 overexpression enhances phagocytosis, activates protective gene programs, and reduces amyloid and neuritic pathology,^13^ while agonist antibodies drive similar effects, improving plaque compaction, lowering Aβ burden, and ameliorating neuronal injury and cognition in vivo.^14,15^

To date, therapeutic strategies aimed at TREM2 have largely focused on antibody-based modalities, while progress in developing small molecule modulators has remained limited. Despite the strong target engagement achieved with antibodies, their clinical translation in AD is hindered by limited blood–brain barrier (BBB) penetration, high cost of manufacture, potential for immunogenic responses, and extended systemic persistence that may prolong adverse events.^16,17^ Small molecule therapeutics provide several advantages in this setting: they generally cross the BBB more effectively, can be delivered orally, offer flexible pharmacokinetic (PK) profiles for dose adjustment, and are less likely to provoke immune reactions.^18^ Their lower production costs also improve the feasibility of widespread patient access.^18^ Incorporating small molecule modulators into immunotherapeutic strategies for AD could therefore enable more versatile and accessible treatments, particularly in combination with other immune-targeting modalities.

Notably, **VG-3927** was recently reported as the first small molecule TREM2 agonist with therapeutic potential in AD, demonstrating strong potency, brain penetration, and disease-modifying effects in preclinical models.^19^ While this finding validates the concept of small molecule TREM2 activation, it also underscores the need for additional chemical scaffolds to diversify therapeutic options and overcome limitations associated with a single chemotype. Motivated by this gap, we pursued a structure-based virtual screening campaign to identify novel small molecule TREM2 modulators. This approach led to the discovery of novel small molecule TREM2 binders, which were subsequently optimized through structure–activity relationship (SAR) studies and validated in functional assays.

To identify chemical starting points for designing TREM2 binders, we have performed an AI-assisted virtual screening workflow (Deep Docking^20^) against the proposed binding site of the TREM2 activator Hecubine (cell-based IC_50_ value of 73 μM^21^), which encompasses parts of both the hydrophobic and basic sites. Briefly, Deep Docking leverages the speed of a deep learning model trained to predict docking scores from small-molecule structure in order to speed up hit discovery. To keep the computational and synthetic lead times as low as possible, we have screened a chemical library of readily purchasable compounds (more than 5 million compounds), following our recent success against challenging PPI-type oncotargets.^22^ The top 30 virtual hits (Table S1) were purchased for bioassay measurements against TREM2, which resulted in one confirmed hit compound, **T2K-014** (Figure 1). Notably, **T2K-014** has a markedly different structure (0.099 Tanimoto similarity of Morgan fingerprints^23^) and interaction profile vs. Hecubine: while **T2K-014** interacts mainly with the hydrophobic site residues W44, L71 and L89 (Figure 1c), the proposed interacting partners of Hecubine are the basic site residues R76, T85 and D87, and the hydrophobic site residue F74 (Figure 1a).^21^ These differences highlight the utility of screening a large chemical space with an AI-assisted docking workflow.

**Figure 1.**
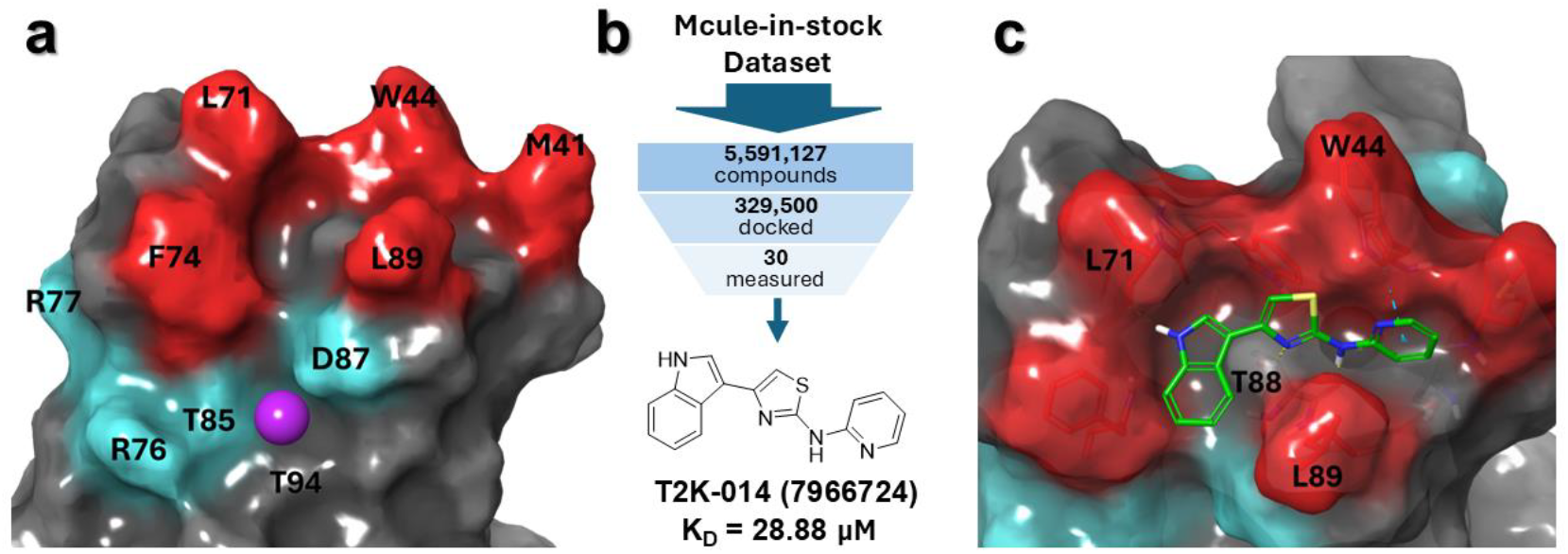
Discovery of TREM2-binding compounds via virtual screening. **(a)** TREM2 binding sites used for virtual screening. Red/cyan residues correspond to the hydrophobic and basic sites, respectively.^24^ The purple sphere represents the proposed binding site of Hecubine.^21^ **(b)** Overview of the virtual screening workflow. With Deep Docking, computational efficiency is maximized by training a deep learning model on a minority of the dataset that is actually docked. **(c)** Predicted binding mode of the confirmed hit **T2K-014**, with the interacting residues labeled. Protein-ligand interactions are shown as dashed lines, with yellow and light blue coloring meaning hydrogen bonds, and pi-pi interactions, respectively.

Experimental binding studies were carried out using the Dianthus platform, which applies Temperature-Related Intensity Change (TRIC) technology to directly monitor protein–ligand interactions. As shown in Figure 2A, candidate compounds were initially screened at 100 μM against purified recombinant human TREM2 protein (10 nM). To validate the assay, we incorporated **PC-192**, a previously described TREM2 agonist, as a positive control, and used protein-only samples to establish baseline variability. Compounds were classified as hits only if their signal exceeded five times the standard deviation of the control.

**Figure 2.**
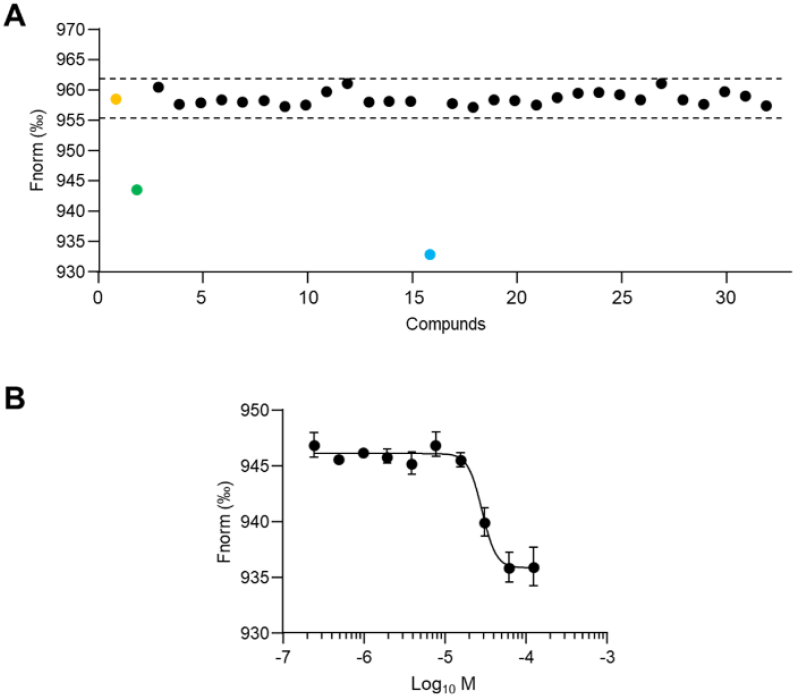
Characterization of selected virtual screening hits for TREM2 binding. **A)** Biophysical validation of 30 computationally selected compounds using TRIC analysis. Compounds were tested at 100 μM against recombinant TREM2 protein (10 nM). Yellow dot: baseline control (TREM2 alone); green dot: reference compound **PC-192**. Dashed lines denote hit selection criteria based on 5-fold standard deviation above background signal. Blue dot represents the confirmed binding compound (**T2K-014**); black dots indicate non-binding compounds. **B)** Dose-response curve using MST for the binding of **T2K-014** to TREM2. Data represent mean ± SEM (n = 3).

From the 30 small molecules examined, **T2K-014** was the only candidate that produced a measurable signal suggestive of binding, although its response was just below the predefined hit threshold, whereas the remainder of the compounds showed no detectable interaction (Figure 2A). This outcome highlights the narrow structural requirements for effective engagement of the TREM2 binding pocket. To further characterize this interaction, we conducted a concentration-dependent analysis using Microscale Thermophoresis (MST). These studies confirmed direct binding of **T2K-014** to TREM2 with a dissociation constant (Kd) of 28.88 ± 1.74 μM (Figure 2B). Although the affinity was modest, this result provided a clear starting point for iterative SAR optimization aimed at improving potency and selectivity.

Guided by the initial virtual screening hit, we pursued a SAR-by-catalog strategy to probe the chemical space around the **T2K-014** scaffold. Twenty-seven commercially available analogs were selected to systematically explore substitutions at two regions of the core structure: the R1 position containing the indole moiety and the R2 position featuring the pyridine group (Table 1). The R1 series included replacements of the indole with alternative heteroaryl or carbocyclic groups, whereas the R2 series introduced variations in ring type and substitution pattern. This design enabled us to contrast the tolerance of the binding pocket to changes at each position, efficiently defining structural determinants of activity and guiding the selection of chemotypes for further optimization. We subsequently conducted primary screening of all 27 derivatives using TRIC analysis at 100 μM concentration against recombinant TREM2 protein (10 nM), with **T2K-014** serving as the reference compound (Figure 3A). From this screening, four compounds (Figure 3B) demonstrated binding signals that exceeded the established threshold criteria. Several compounds exhibited aggregation behavior or interference effects that prevented reliable binding assessment.

**Table 1.**
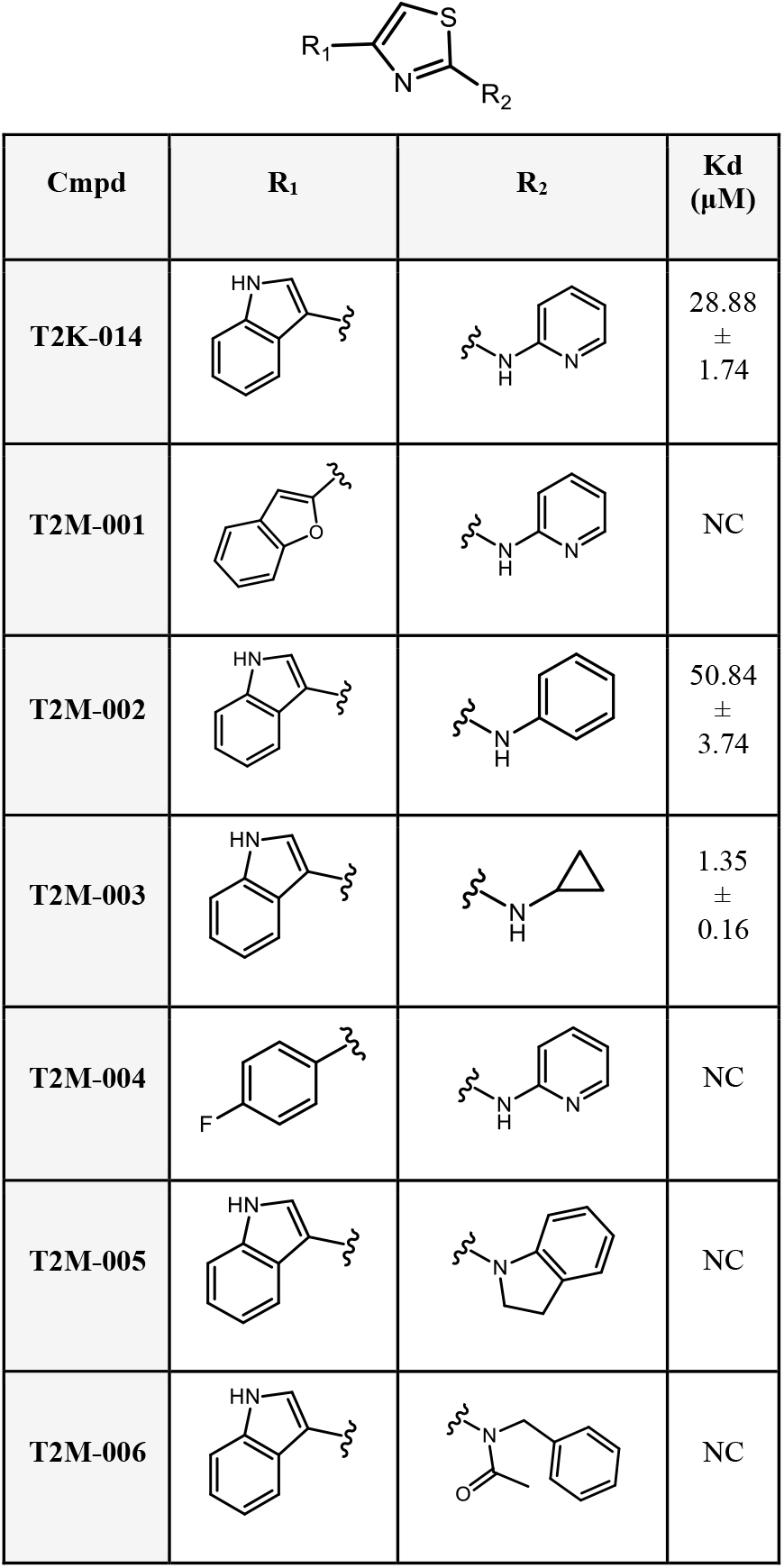
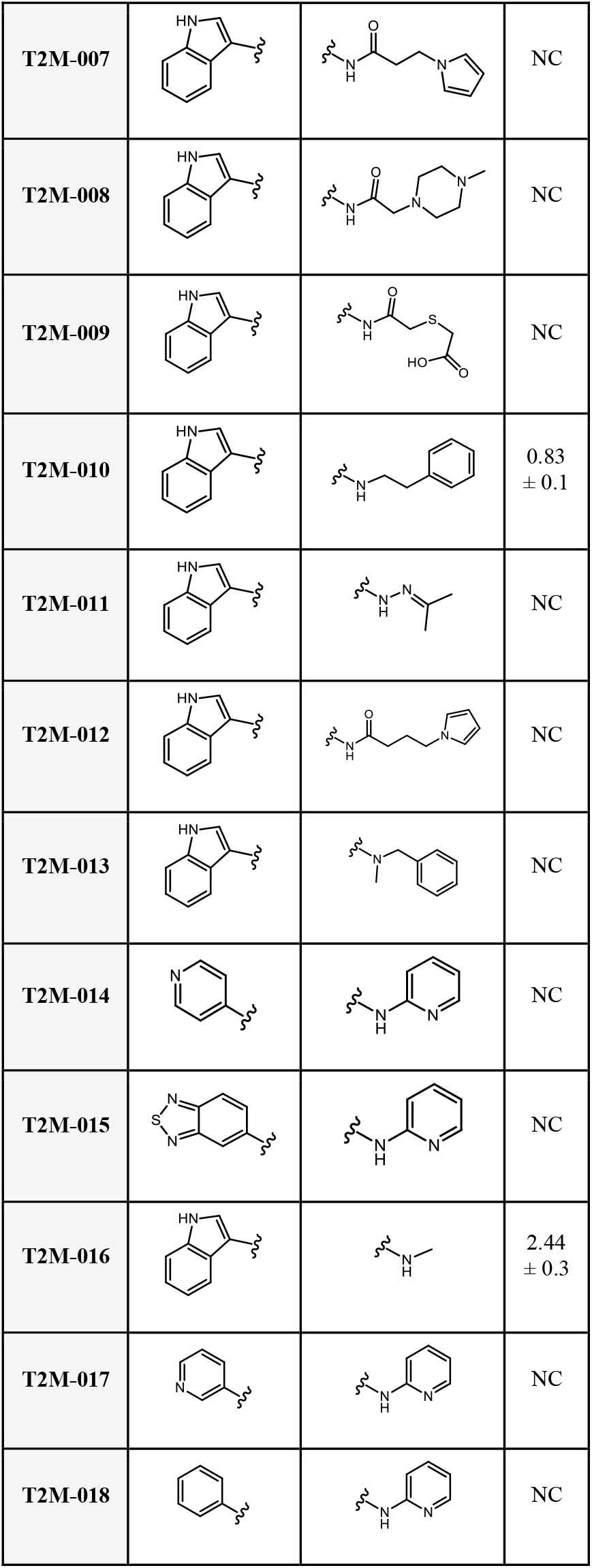
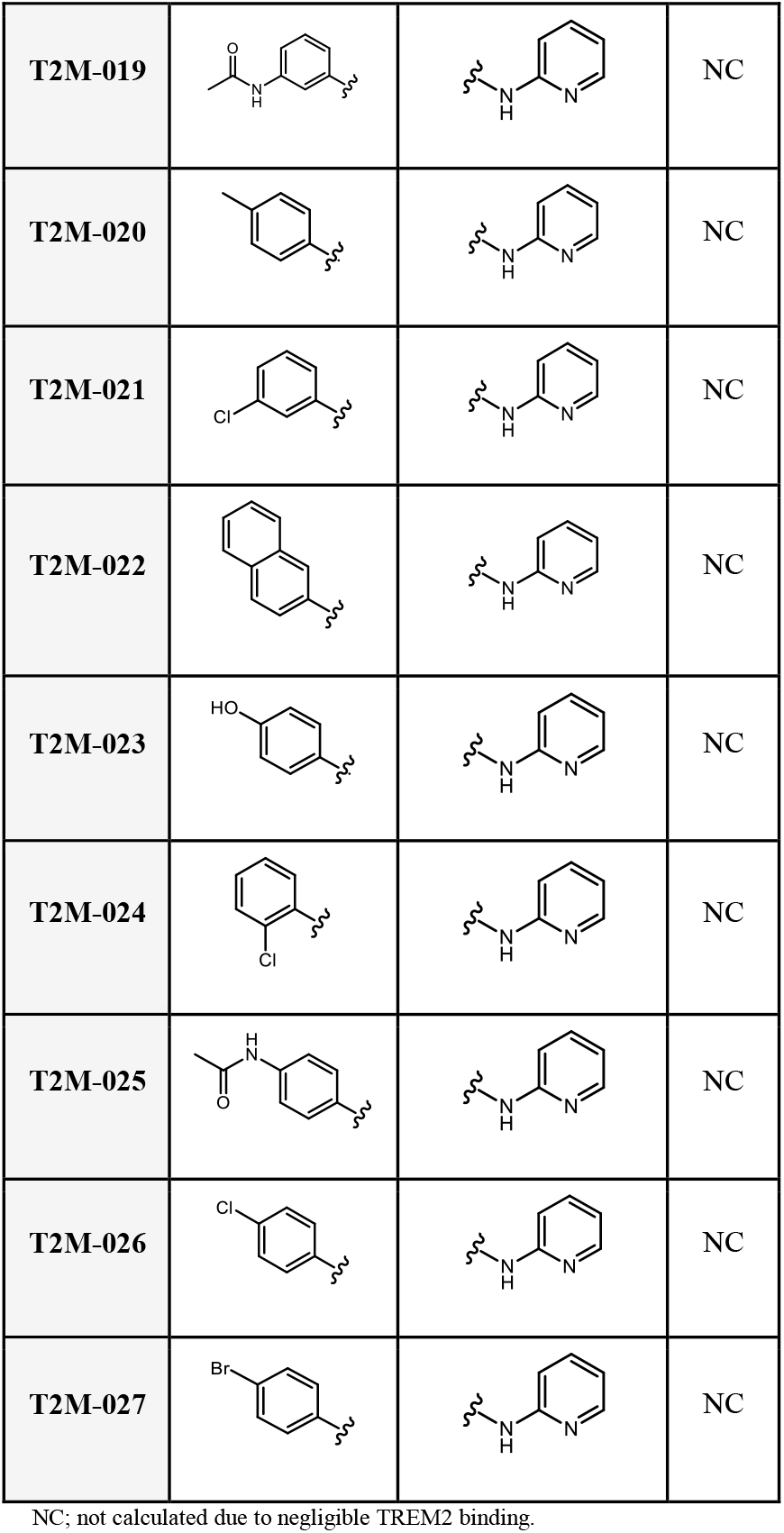
Chemical structures and TREM2 binding affinity (Kd) of T2K-014 derivatives.

**Figure 3.**
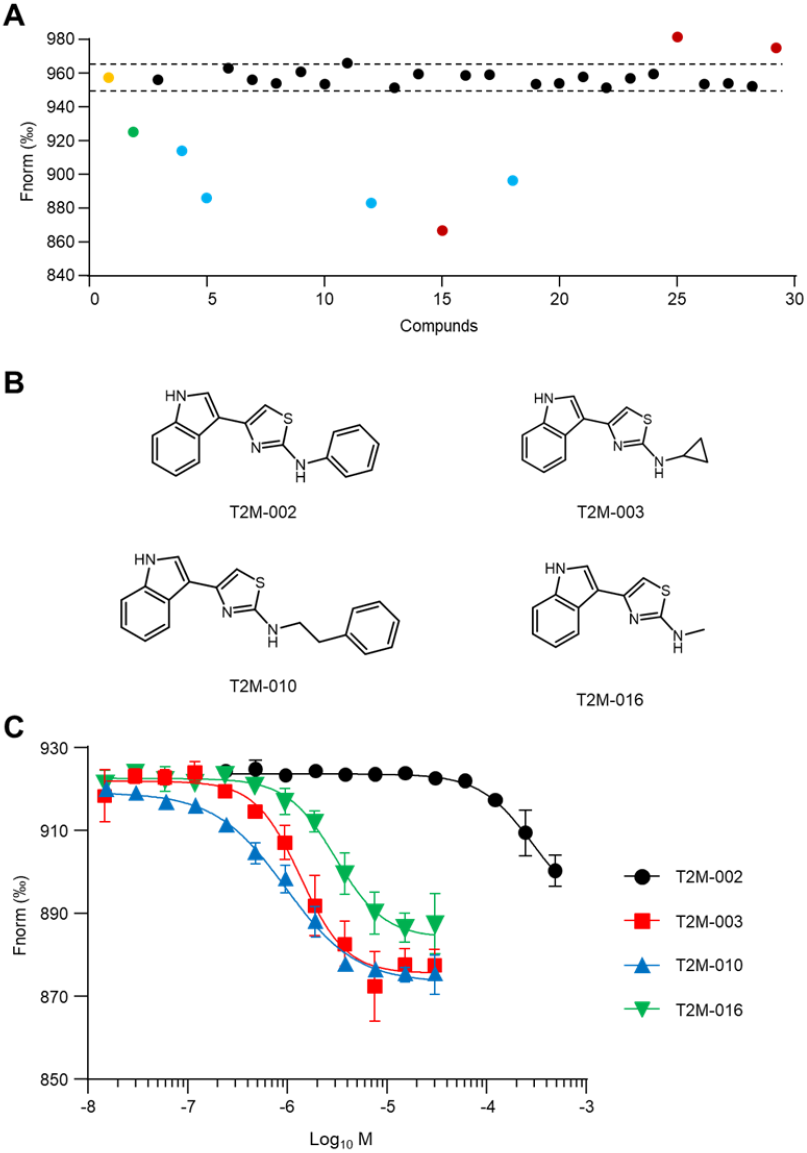
SAR-by-catalog study of T2K-014 derivatives. **A)** Single-concentration screening of 27 **T2K-014** derivatives against TREM2 protein (10 nM) at 100 μM using TRIC analysis. Yellow dot: negative control (TREM2 alone); green dot: positive control (**T2K-014**). Dashed lines represent the threshold for hit identification, corresponding to signals exceeding 5-fold the standard deviation of the negative control. Blue dots indicate validated TREM2-binding hits; red dots represent compounds causing aggregation or fluorescence interference. **B)** Chemical structures of four **T2K-014** derivatives identified through TRIC screening: **T2M-002, T2M-003, T2M-010**, and **T2M-016. C)** Concentration-dependent binding profiles of **T2K-014** derivatives determined by MST. Black circles: **T2M-002**; red squares: **T2M-003**; blue triangles: **T2M-010**; green inverted triangles: **T2M-016**. Data are presented as mean ± SEM (n = 3).

As shown in Figure 3C, the four hits (**T2M-002, T2M-003, T2M-010**, and **T2M-016**) were evaluated to quantitative binding analysis using MST to determine their TREM2 binding affinities. **T2M-002** displayed a Kd value of 50.84 ± 3.74 μM, representing slightly weaker binding than the parent compound (**T2K-014**). However, three derivatives demonstrated improved affinities: **T2M-003** achieved a Kd of 1.35 ± 0.16 μM, **T2M-016** exhibited a Kd of 2.44 ± 0.30 μM, and **T2M-010** showed the strongest binding with a Kd of 0.83 ± 0.10 μM. These improvements represent approximately 21-fold, 12-fold, and 35-fold enhancements in binding affinity, respectively, compared to **T2K-014**. To the best of our knowledge, **T2M-010** is the most potent small molecule TREM2 binder reported to date.

Structural analysis of the active analogs revealed a clear trend: all four hits preserved the indole moiety at the R1 position, whereas modifications at this site were uniformly detrimental to activity (Table 1). Although this conclusion is based on a limited set of variants, it points to the indole as a critical pharmacophore element for TREM2 engagement. In contrast, diversity at the R2 position was more compatible with binding, suggesting that this region offers chemical flexibility for tuning potency and selectivity. Within this series, **T2M-010** emerged as the most promising analog, establishing it as an optimized lead and highlighting the R2 site as a productive avenue for continued medicinal chemistry exploration.

The functional studies provide compelling evidence that **T2M-010** engages TREM2 in a biologically meaningful manner. The phosphorylation assays established that **T2M-010** is not only a binder but also a functional activator of TREM2 signaling. In HEK–hTREM2/DAP12 cells, treatment with **T2M-010** led to a robust twofold increase in SYK phosphorylation, a canonical downstream readout of TREM2 activation, comparable in magnitude to the benchmark TREM2 agonist **VG-3927** (Figure 4). This finding is critical because it demonstrates that the structural optimization achieved during SAR efforts translated into a compound capable of eliciting receptor-mediated signaling. By contrast, **T2M-016** induced only a modest, non-significant change, suggesting that small variations in scaffold design strongly influence signaling competence. These results reinforce the conclusion that **T2M-010** occupies the binding pocket in a productive orientation that effectively couples receptor engagement to downstream signal transduction. Establishing this link between biochemical binding and cellular signaling represents a key step in validating **T2M-010** as a TREM2 agonist rather than a non-functional binder.

**Figure 4.**
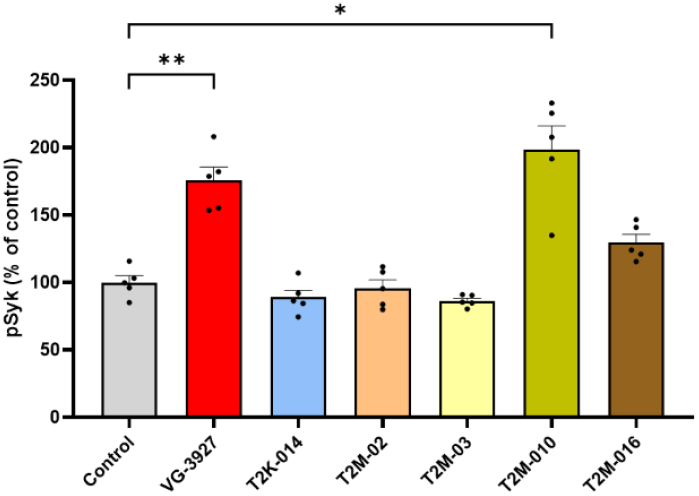
Relative changes in SYK phosphorylation in response to TREM2 agonists, measured by AlphaLISA. Histogram illustrating the relative levels of phosphorylated SYK (Tyr525/526) in HEK–hTREM2/DAP12 cells under control conditions and following treatment with candidate TREM2 agonists: **T2K-014, T2M-02, T2M-03, T2M-010** and **T2M-016** at 25 µM for 1 hour. Additionally, cells were treated with the commercial TREM2 agonist **VG-3927** (25 µM, 1 h) as a reference compound. Phospho-SYK levels were quantified using the AlphaLISA SureFire Ultra assay and are expressed as a percentage relative to the control condition (vehicle-treated cells). Data represent the mean ± SEM from five independent biological replicates. Statistical significance was assessed using Brown-Forsythe and Welch Anova test followed by Dunnett ‘s multiple comparison test (\**p* <0.05, \*\**p* < 0.01).

Beyond proximal signaling events, we next assessed whether **T2M-010** could drive a functional microglial phenotype. In BV2 cells, **T2M-010** significantly enhanced bead uptake, demonstrating a marked increase in phagocytic activity that paralleled the effect of **VG-3927** (Figure 5). This is a critical functional endpoint, as microglial phagocytosis is central to the clearance of amyloid and cellular debris in AD models. The ability of **T2M-010** to stimulate this process underscores its capacity to recapitulate protective microglial programs previously linked to genetic or antibody-mediated TREM2 activation. Importantly, the consistency between the SYK phosphorylation and phagocytosis assays establishes a coherent mechanism of action, in which TREM2 binding triggers intracellular signaling cascades that culminate in enhanced microglial effector function. Together, these results place **T2M-010** within a class of small molecules capable of promoting microglial responses relevant to disease modification.

**Figure 5.**
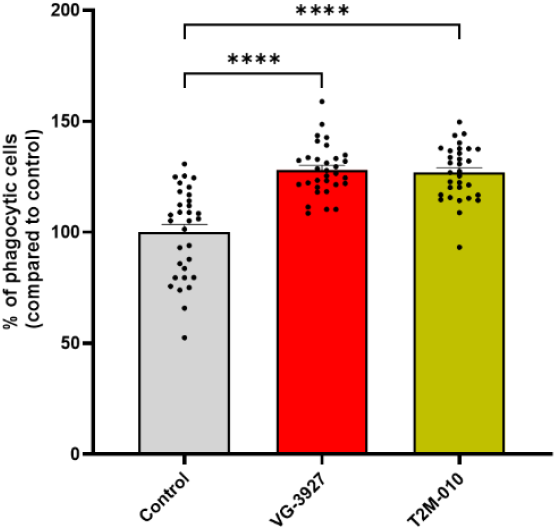
Quantification of phagocytic activity in BV2 microglial cells. BV2 cells were pre-treated with the indicated compounds (**T2M-010** or **VG-3927**) at a concentration of 25 µM, or vehicle control (DMSO) for 30 minutes, followed by exposure to green fluorescent latex beads. Phagocytic cells were defined as IBA1-positive cells containing at least one internalized fluorescent bead. Quantification is expressed as the percentage of phagocytic cells relative to vehicle control. Data represent mean ± SEM from 32 images per group across four independent biological replicates (n = 4). Statistical significance was assessed using Brown-Forsythe and Welch ANOVA followed by Dunnett’s multiple comparisons test. \*\*\*\**p* < 0.0001.

To gauge the CNS drug-likeness of **T2M-010**, we completed a focused *in vitro* ADME/PK assessment (Table 2). The compound exhibits a balanced physicochemical profile, with a moderate logD_7.4_ of 2.8 and kinetic solubility of 48 µM in PBS (1% DMSO). Its FaSSIF solubility of 95 µM suggests adequate dissolution under fasted intestinal conditions and supports oral absorption potential. Indicators of BBB passage were favorable (Table 2). In PAMPA-BBB, **T2M-010** showed high passive permeability (P_e_ = 9.2 × 10^−6^ cm/s). Consistently, MDCK– MDR1 assays returned robust A→B transport (Papp = 2.85 × 10^−5^ cm/s) and a low efflux ratio (ER = 1.07), pointing to minimal P-gp–mediated efflux (Table 2). Together, these data indicate that passive diffusion is likely the dominant driver of CNS exposure for **T2M-010**, a profile compatible with efficient brain penetration.

**Table 2.**
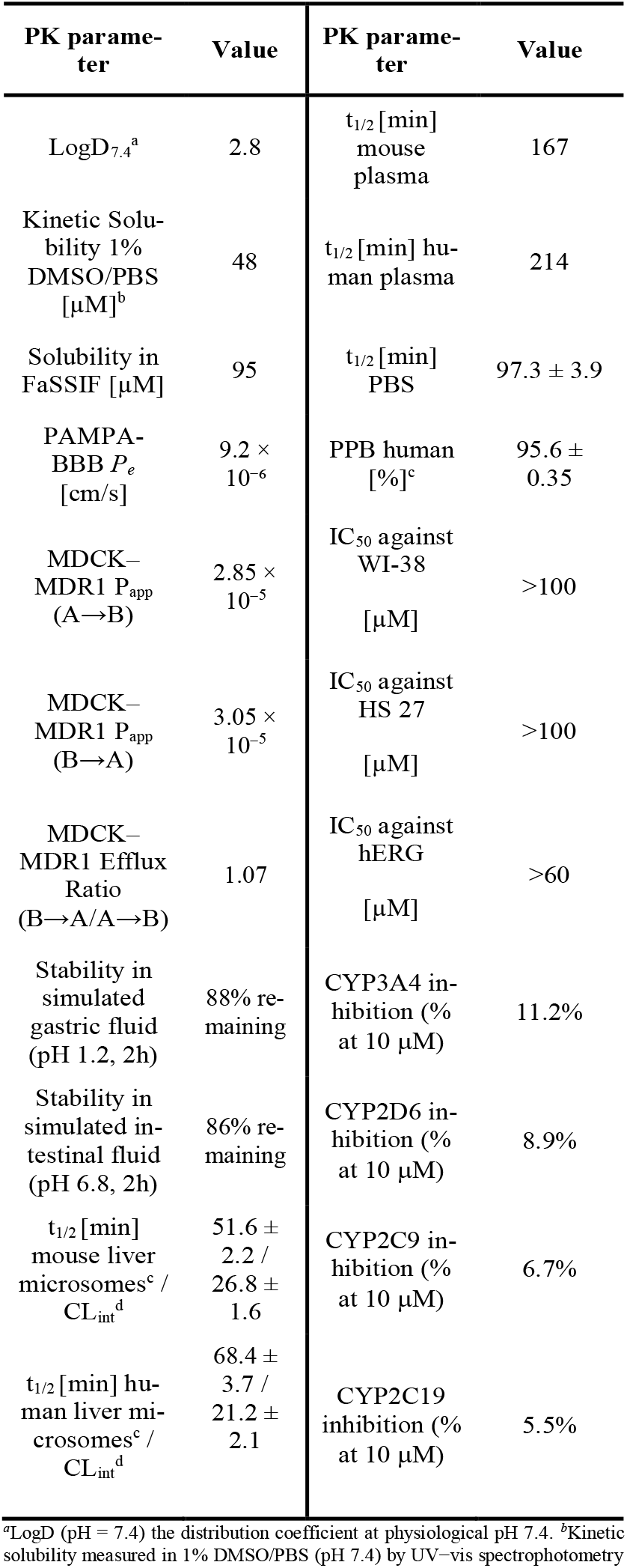
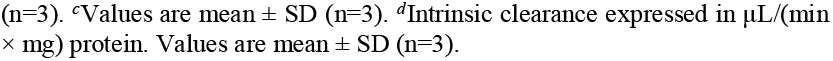
In vitro PK profile of **T2M-010**.

**T2M-010** also displayed good gastrointestinal stability (>88–86% parent remaining after 2 h in simulated gastric and intestinal fluids, respectively) and long plasma half-lives (167 min in mouse; 214 min in human). The compound was chemically stable in buffer (t_1/2_ = 97.3 ± 3.9 min in PBS). Microsomal turnover was moderate, with mouse and human liver microsome half-lives of 51.6 ± 2.2 min (CL_int_ = 26.8 ± 1.6 μL/min/mg) and 68.4 ± 3.1 min (CL_int_ = 21.2 ± 2.1 μL/min/mg), respectively-consistent with an acceptable clearance projection for a CNS small molecule (Table 2). In safety pharmacology counterscreens, **T2M-010** showed no cytotoxicity in human fibroblasts (WI-38 and HS-27; IC_50_ >100 µM), low hERG risk (IC_50_ >60 µM), and minimal inhibition of major CYP isoforms at 10 µM (≤11.2%). Overall, the ADME/PK profile of **T2M-010** (Table 2) supports continued preclinical progression, balancing solubility, BBB permeability, metabolic stability, and preliminary safety.

In this study, we demonstrate that an AI-assisted virtual screening approach can successfully identify small molecule modulators of TREM2, a receptor of central importance to microglial biology and AD pathogenesis. Starting from a modest-affinity hit, **T2K-014**, a SAR-by-catalog campaign revealed key structural requirements for activity and yielded **T2M-010** as an optimized analog with submicromolar binding affinity. **T2M-010** not only displayed favorable in vitro ADME/PK properties consistent with CNS drug-likeness but also engaged TREM2 in a functionally meaningful manner, inducing SYK phosphorylation and enhancing microglial phagocytosis to levels comparable with a benchmark agonist. These findings establish **T2M-010** as the most potent small-molecule TREM2 binder reported to date and provide proof-of-concept that small molecules can reproduce protective microglial programs previously accessible only through antibody-based strategies. While further medicinal chemistry and in vivo validation are essential, the present work delivers a validated chemical scaffold and a clear roadmap toward the development of orally bioavailable TREM2 agonists as potential therapeutics for AD.

## Supporting Information

The Supporting Information is available free of charge on the ACS Publications website.

Computational methods, biophysical screening assays, cellular evaluation, and chemical structures of selected compounds (PDF)

## Author information

### Author Contributions

The manuscript was written through contributions of all authors. All authors have given approval to the final version of the manuscript.

## Abbreviations

AD: Alzheimer’s disease
Aβ: amyloid-β
BBB: blood–brain barrier
CNS: central nervous system
ER: efflux ratio
FaSSIF: fasted state simulated intestinal fluid
IC_50_: half-maximal inhibitory concentration
ITAM: immunoreceptor tyrosine-based activation motif
Kd: equilibrium dissociation constant
MST: microscale thermophoresis
NC: not calculated
PBS: phosphate-buffered saline
PK: pharmacokinetics
SAR: structure–activity relationship
SYK: spleen tyrosine kinase
TREM2: triggering receptor expressed on myeloid cells 2
TRIC: Temperature-Related Intensity Change.

## Notes

The authors declare no competing financial interests.

## Acknowledgments

This work was supported by the National Institutes on Aging under grant number R01AG083512 (PI: Gabr). This work was supported by the National Research Development and Innovation Office of Hungary [contracts FK146063 to D.B., Pharma-Lab (RRF-2.3.1-21-2022-00015) to G.M.K.]. The work of D.B. was supported by the János Bolyai Research Scholarship of the Hungarian Academy of Sciences.

## Insert Table of Contents artwork here

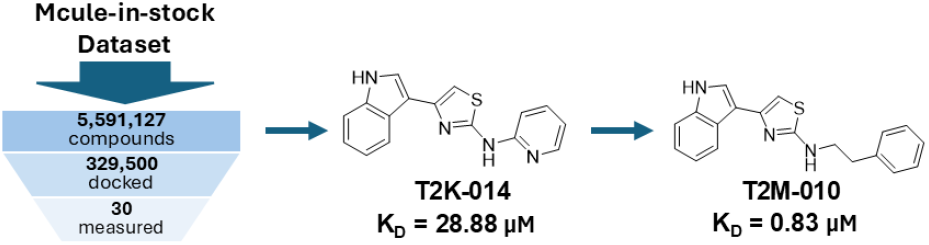

