## Supporting Information for "AI-Assisted Discovery and Optimization of Small Molecule TREM2 Agonists with Functional Microglial Activity"

*Electronic Supplementary Information*

| **Contents** |  |
| --- | --- |
| Experimental procedures | S2 |
| Chemical structures of virtual screening hits as potential TREM2-targted small molecules  References | S5  S7 |

**Experimental procedures**

**1. Docking and virtual screening**

The protein structure with the PDB ID 6Y6C^1^ was used for docking. The structure was prepared using Schrödinger Maestro’s Protein Preparation Workflow^2^ at a pH of 7.4. The receptor grid was set by including the binding site of the known small-molecule activator Hecubine, the hydrophobic site and a part of the basic site.^3-5^

For ligands, the Mcule-in-stock library for readily purchasable compounds (accessed at 2022. 11. 04.) with 5,591,127 compounds were used.^6^ PAINS compounds were filtered out from the dataset in advance.^7^

Virtual screening was performed using the AI-based tool Deep Docking^1^ with a total of eleven iterations, each with 21,500 ligands to be docked (3 × 21,500 in the first iteration). Other Deep Docking parameters (e.g. number of hyperparameters, recall value) were set to the default values as described in the original paper.^8^ In each iteration, LigPrep^2^ was used to prepare the ligands at a pH range of 7.4±1.0, and Glide SP^9,10^ was used as the docking algorithm. In total, 279,500 compounds were used for Deep Docking, which corresponds to about ~5% of the whole dataset. After the last iteration, for cherry-picking purposes, 50,000 compounds were exported using the trained model, which were prepared in the same way as described earlier, and were docked using Glide SP.

To enhance chemical diversity, the Score Erosion node^11^ in KNIME^12^ was used with an erosion factor of 0.2 to deprioritize chemically similar compounds. In total, 30 compounds were chosen for experimental testing based on docking scores, chemical structure (heavy atoms ≥ 15, molar mass ≤ 700 Da, amide bonds ≤ 3) and binding mode (at least three advantageous interactions with the protein and no significant solvent-exposed part).

**2. Temperature-Related Intensity Change (TRIC) Binding Assay**

Temperature-Related Intensity Change (TRIC) measurements were performed using a Dianthus NT.23PicoDuo instrument (NanoTemper Technologies, Munich, Germany). Recombinant human TREM2 protein (in-house) was labeled with RED-tris-NTA 2nd Generation dye using the His-Tag Labeling Kit (NanoTemper Technologies) according to the manufacturer's protocol. All binding experiments were conducted in PBST buffer containing 154 mM NaCl, 5.6 mM Na₂HPO₄, 1.05 mM KH₂PO₄, pH 7.4, and 0.005% Tween-20. For binding measurements, labeled TREM2 protein was used at a final concentration of 10 nM. Test compounds were serially diluted in PBST buffer and incubated with labeled TREM2 for 10 minutes at room temperature (22-25°C) prior to analysis. The Dianthus instrument was configured with the following parameters: 85% LED excitation power, picomolar detector sensitivity disabled, and laser on-time of 5 seconds. Data analysis was performed using Dianthus Analysis software (NanoTemper Technologies) for initial processing, followed by curve fitting and statistical analysis using GraphPad Prism 10.0 (GraphPad Software, San Diego, CA, USA).

**3. Microscale Thermophoresis (MST) Binding Assay**

Binding affinity measurements were performed using microscale thermophoresis (MST) on a Monolith NT.115 system (NanoTemper Technologies, Munich, Germany). Recombinant human TREM2 protein (in-house) and human TREM1 protein (BioTechne, Minneapolis, MN, USA) were labeled using the RED-tris-NTA His-tag labeling kit (NanoTemper Technologies) according to the manufacturer's instructions. MST measurements were conducted using different buffer conditions optimized for each protein: TREM2 assays were performed in PBS buffer (pH 7.4) containing 0.005% Tween-20, while TREM1 assays used PBS buffer (pH 7.4) with 0.05% Tween-20. Labeled proteins were used at a final concentration of 40 nM and incubated with serially diluted test compounds for 10 minutes at room temperature (22-25°C) prior to measurement. MST experiments were performed using standard capillaries with the following instrument parameters: red filter set, 100% LED power, and medium MST power. Thermophoresis was monitored for 20 seconds with an additional 5-second delay. Data analysis was conducted using MO.Affinity Analysis software (NanoTemper Technologies) for initial processing, followed by curve fitting using GraphPad Prism 10.0 (GraphPad Software, San Diego, CA, USA).

**4. Cell Culture and Treatment**

HEK–hTREM2/DAP12 cells were seeded at a density of 5 × 10⁴ cells per well in a 96-well plate containing 100 µL of Dulbecco’s Modified Eagle Medium (DMEM; Gibco), supplemented with 10% fetal bovine serum (FBS; Gibco). Cells were incubated for 24 hours at 37 °C in a humidified atmosphere with 5% CO₂. Subsequently, the growth medium was replaced with fresh medium containing test compounds at a final concentration of 25 µM. Vehicle control wells received an equivalent volume of dimethyl sulfoxide (DMSO), ensuring a final concentration of 0.25%. Cells were incubated for an additional hour under the same conditions.

**5. Phospho-SYK Detection**

After treatment, culture medium was carefully removed, and cells were lysed directly in the wells using the lysis buffer provided by the supplier (Revvity, USA) and pSYK (Tyr525/526) was measured using AlphaLisa technique according to the manufacturer’s protocol. This assay employs two specific antibodies: one targeting phosphorylated Tyr525/526 on SYK, and another binding a distinct epitope on SYK. When phospho-SYK is present, donor and acceptor beads brought into proximity by these antibodies facilitate singlet oxygen transfer, resulting in a luminescent Alpha signal. The intensity of luminescence is directly proportional to the phospho-SYK concentration in the sample. Assay plate included a positive control (phospho-SYK-enriched cell lysate supplied with the kit) and a negative control (lysis buffer only). All samples and controls were processed concurrently to maintain assay consistency and reliability.

**6. Phagocytosis Assay in BV2 Microglial Cells**

Phagocytic activity of BV2 microglial cells was evaluated using green fluorescent latex beads as previously described.^13^ Briefly, cells treated for 30min with either 25 µM T2M-010, vehicle control (DMSO) or VG-3927 that served as a reference. Following compound exposure, fluorescent beads were added to the culture medium, and after incubation, cells were processed for immunocytochemistry. Imaging was performed using a fluorescence microscope, DAPI was used for nuclear counterstaining, and IBA1 immunolabeling identified microglial cells. Cells were considered phagocytically active if they were IBA1-positive and contained at least one internalized fluorescent bead.

**7. Immunocytochemistry**

Fixed cells were permeabilized with PBS containing 0.25% Triton X-100 and blocked with 1% bovine serum albumin (BSA). Cells were then incubated with primary anti-IBA1 antibody, followed by a secondary Alexa Fluor 594-conjugated antibody.

**8. Pharmacokinetics (PK) study**

The preliminary evaluation of PK parameters for As48 was performed as previously reported by us.^13^ In brief, physicochemical and biological profiling included determination of LogD₇.₄, microsomal stability, kinetic solubility, and cytotoxicity across a panel of cell lines. Solubility was assessed using UV–vis spectrophotometry, while cell viability was measured using the PrestoBlue® assay (ThermoFisher, Cat# A13261).

**Table S1. Chemical structures of virtual screening hits as potential TREM2-targted small molecules.**

| 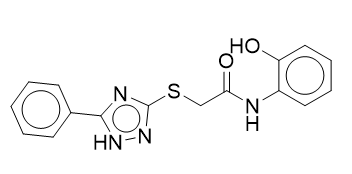 | 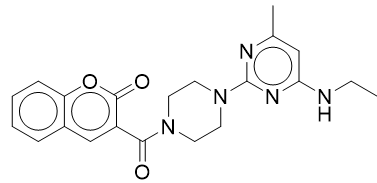 |
| --- | --- |
| 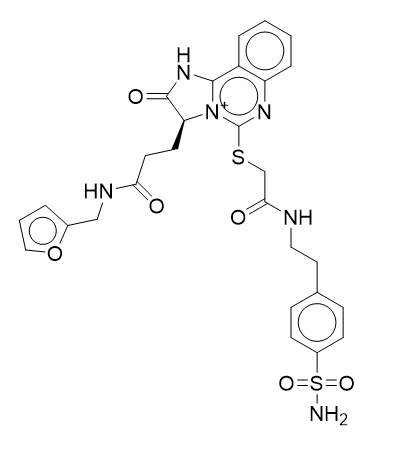 | 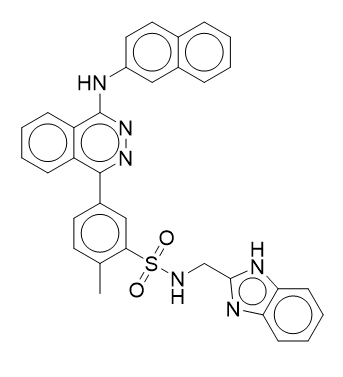 |
| 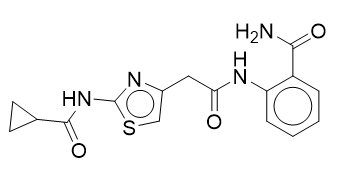 | 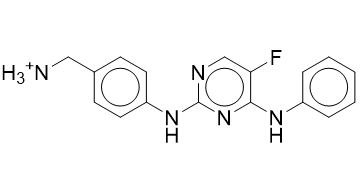 |
| 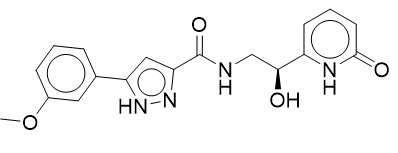 | 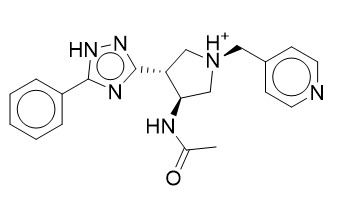 |
| 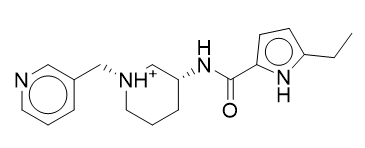 | 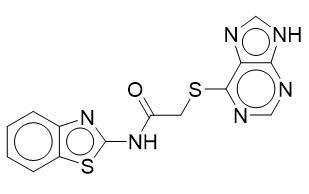 |
| 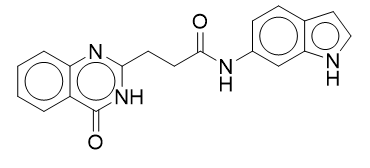 | 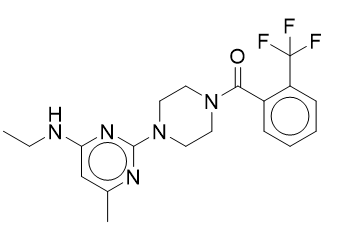 |
| 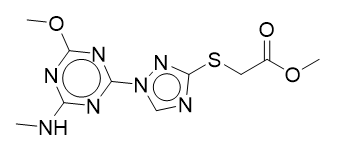 | 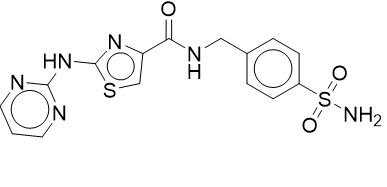 |
| 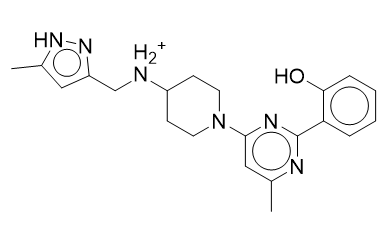 | 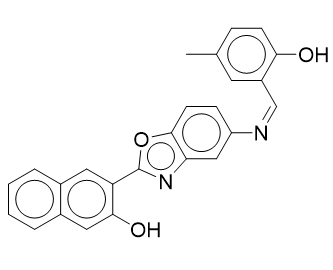 |
| 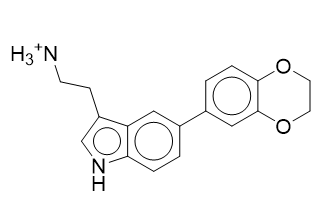 | 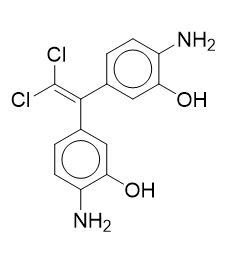 |
| 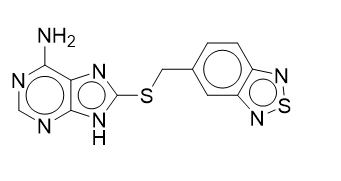 | 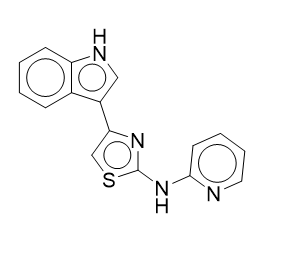 |
| 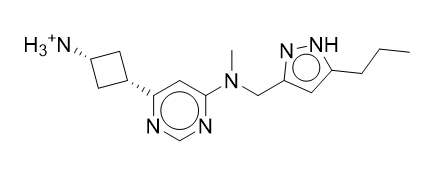 | 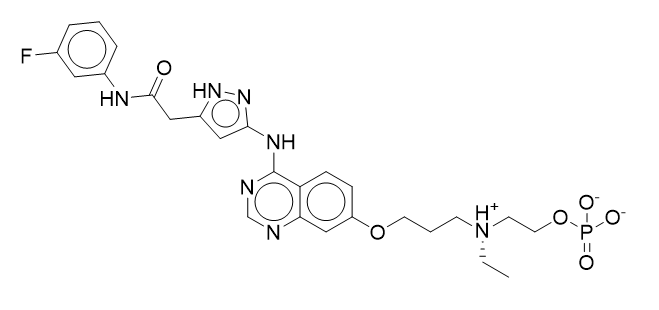 |
| 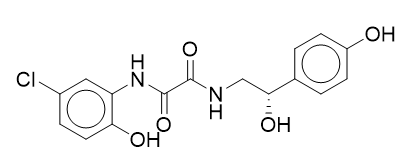 | 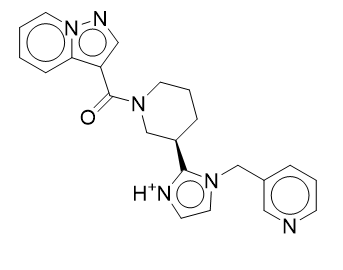 |
| 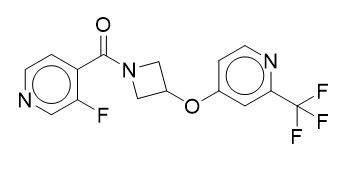 | 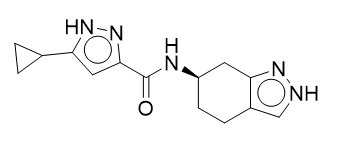 |
| 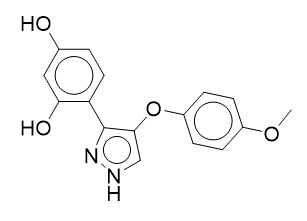 | 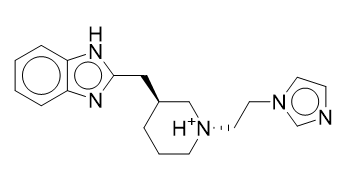 |
| 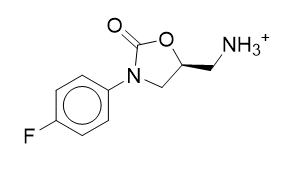 | 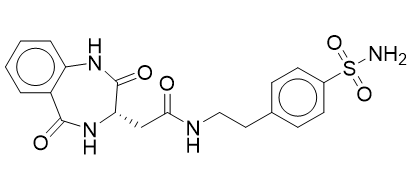 |
